# *Listeria* adhesion protein orchestrates caveolae-mediated apical junctional remodeling of epithelial barrier for *L. monocytogenes* translocation

**DOI:** 10.1101/2023.10.18.563002

**Authors:** Rishi Drolia, Shivendra Tenguria, Donald B. Bryant, Jessie Thind, Breanna Amelunke, Dongqi Liu, Nicholas L.F. Gallina, Krishna K. Mishra, Manalee Samaddar, Manoj R. Sawale, Dharmendra K. Mishra, Abigail Cox, Arun K Bhunia

**Affiliations:** Molecular Food Microbiology Laboratory, Department of Food Science, Purdue University, West Lafayette, Indiana, USA; Department of Biological Science, Old Dominion University, Norfolk, Virginia, USA; Department of Biological Science, Eastern Kentucky University, Richmond, Kentucky 40475, USA; Purdue Institute of Inflammation, Immunology, and Infectious Disease, Purdue University, West Lafayette, Indiana, USA; Department of Comparative Pathobiology, Purdue University, West Lafayette, Indiana, USA

**Keywords:** *Listeria monocytogenes*, *Listeria* Adhesion Protein (LAP), Heat Shock Protein 60 (Hsp60), Tight Junction (TJ), Intestinal Barrier, Internalin A (InlA), Translocation, E-cadherin, Caveolin, Dynamin

## Abstract

The cellular junctional architecture remodeling by LAP-Hsp60 interaction for *L. monocytogenes* (*Lm*) passage through the epithelial barrier is incompletely understood. Here, using the gerbil model, permissive to internalin (Inl) A/B-mediated pathways like in humans, we demonstrate that *Lm* crosses the intestinal villi at 48 h post-infection. In contrast, the single isogenic (*lap^─^* or Δ*inlA*) or double (*lap^─^ΔinlA*) mutant strains show significant defects. LAP promotes *Lm* translocation via endocytosis of cell-cell junctional complex in enterocytes that do not display luminal E-cadherin. In comparison, InlA-mediated transcytosis occurs in enterocytes displaying apical E-cadherin during cell extrusion and mucus expulsion from goblet cells. LAP hijacks caveolar endocytosis to traffic integral junctional proteins to the early and recycling endosomes. Pharmacological inhibition in a cell line and genetic knock-out of caveolin-1 in mice prevents LAP-induced intestinal permeability, junctional endocytosis, and *Lm* translocation. Furthermore, LAP-Hsp60-dependent tight junction remodeling is also necessary for InlA access to E-cadherin for *Lm* intestinal barrier crossing in InlA-permissive hosts.

## IMPORTANCE

*Listeria monocytogenes* (*Lm*) is a foodborne pathogen with high mortality (20-30%) and hospitalization rates (94%), particularly affecting vulnerable groups such as pregnant women, fetuses, newborns, seniors, and immunocompromised individuals. Invasive listeriosis involves *Lm’s* internalin (Inl) A protein binding to E-cadherin to breach the intestinal barrier. However, non-functional InlA variants have been identified in *Lm* isolates, suggesting InlA-independent pathways for translocation. Our study reveals that *Listeria* adhesion protein (LAP) and InlA cooperatively assist *Lm* entry into the gut lamina propria in a gerbil model, mimicking human listeriosis in early infection stages. LAP triggers caveolin-1-mediated endocytosis of critical junctional proteins, transporting them to early and recycling endosomes, facilitating *Lm* passage through enterocytes. Furthermore, LAP-Hsp60-mediated junctional protein endocytosis precedes InlA’s interaction with basolateral E-cadherin, emphasizing LAP and InlA’s cooperation in enhancing *Lm* intestinal translocation. This understanding is vital in combating the severe consequences of *Lm* infection, including sepsis, meningitis, encephalitis, and brain abscess.

## INTRODUCTION

The mucosa of the intestines is lined by epithelial cells that provide the first line of defense against infection. The intestinal barrier comprises a single layer of polarized columnar epithelial cells (enterocytes) organized into finger-like projections called villi (1–3). These cells are self-renewed every 4–5 days, making the intestinal epithelium a highly dynamic structure. The intestinal epithelial cells are polarized, with the apical surface facing the lumen and the basal surface facing the basement membrane. Epithelial cells are networked through adhesive contacts called junctions, which join cells and provide a paracellular seal. The paracellular space between two adjacent epithelial cells from the apical to basolateral direction is sealed by tight junctions (TJs), and adherens junctions (AJs) comprise the apical junctional complex (AJC) that tightly regulate the intestine’s barrier function (4–6). The endocytic pathways involved in junctional regulation are crucial for the dynamic maintenance of cell-cell adhesion and the permeability of epithelial and endothelial barriers (7). In compromised conditions, the AJC can be endocytosed, resulting in the breach of the barrier, thus allowing the invasion of pathogens into the underlying sterile tissue (8). The precise mechanism of how entero-invasive pathogens remodel the cellular architecture and breach the AJC to cause systemic infection remains poorly defined.

*Lm* is a model facultative intracellular and an opportunistic foodborne pathogen. In high-risk populations, neonates, the aged, and immunocompromised individuals, it can cause meningitis, encephalitis, liver abscessation and stillbirth, and abortion in pregnant women with a case fatality rate of 20-30% (9,10). The bacterium can efficiently cross host barriers, including intestinal, blood-brain, and placental (1–3). The intestinal tract is the primary route for *Lm* infection, and crossing this barrier is a crucial first step before spreading to deeper tissues (11).

The *Lm* invasion proteins, internalin A (InlA) and internalin B (InlB), play a pivotal role in internalizing *Lm* into human nonphagocytic cells. However, the interaction of both InlA and InlB with their respective receptors is host species-specific (12). InlA interacts with human and guinea pig E-cadherin to promote bacterial transcytosis across the gut epithelium (13,14) but not with mouse or rat E-cadherin (15,16). InlB interacts with human and mouse Met to accelerate *L. monocytogenes* invasion of the Peyer’s patches (17,18), but not with a guinea pig or rabbit Met (19). In contrast, the gerbil, like humans, is the only small animal model that is naturally permissive to both InlA/E-cadherin and InlB/Met interactions, similar to humans (20).

E-cadherin at the adherens junction (AJ) is inaccessible to the luminal *Lm* InlA. Host intrinsic mechanisms such as epithelial cell extrusion at the tip of the villi (21,22) and mucus exocytosis from goblet cells (13,23) allow E-cadherin to be transiently luminally accessible, facilitating transcytosis across the intestinal barrier without spreading from cell-to-cell. However, it is unclear whether *Lm* actively induces alterations of the AJC to access basolateral E-cadherin for *Lm* translocation. While the importance of InlA-E-cadherin interaction in the bacterial crossing of the intestinal epithelial barrier is critical (13,16,20), *Lm* expressing a non-functional truncated InlA have been isolated from patients (24,25) and fetuses of pregnant guinea pigs after oral dosing (26,27). These suggest that the InlA-independent mechanism(s) is critical for *Lm* translocation across the intestinal barrier and relevant to human infection.

Our group has demonstrated the crucial role of *<u>L</u>isteria* <u>a</u>dhesion <u>p</u>rotein (LAP) as an adhesion molecule in *Lm* pathogenesis during the intestinal phase of infection (28,29). LAP is a 94-kDa alcohol acetaldehyde dehydrogenase (AdhE; *lmo1634*) is a moonlighting protein that promotes bacterial adhesion to cell lines of intestinal origin only in pathogenic *Listeria* species (11,30,31). LAP is a SecA2-dependent secreted protein (32,33) anchored to InlB on the *Lm* cell wall (34). We also deciphered the human heat shock protein 60 (Hsp60) as the host epithelial receptor for LAP (11,35,36). LAP promotes bacterial translocation into the lamina propria and systemic dissemination by increasing intestinal epithelial permeability *in vivo* in mice (InlA-non permissive) (6). By interacting with its cognate receptor, Hsp60, LAP activates canonical NF-κB(p65) signaling, thereby promoting the myosin light chain kinase (MLCK)-mediated opening of the intestinal cell-cell junction via the cellular redistribution of the major junctional proteins and promoting bacterial translocation (6). However, how LAP orchestrates *Lm* translocation in a human-relevant InlA-permissive host and the precise mechanism of junctional remodeling *in vivo* are not established.

Here, we show that both LAP and InlA promote *Lm* translocation into the lamina propria in a human-relevant InlA/E-cadherin and InlB/Met permissive gerbil model during the early stage of infection. Mechanistically, the *Lm* LAP initiates caveolin-1, mediates endocytosis of integral apical junctional proteins, and traffics them to the early and recycling endosomal intracellular destinations for *Lm* translocation across enterocytes. We further demonstrate that LAP-Hsp60-mediated junctional protein endocytosis is critical for providing InlA access to basolateral E-cadherin. Our findings suggest LAP and InlA cooperatively promote *Lm* translocation across the intestinal epithelial barrier for successful infection.

## RESULTS

### *Lm* translocate across the intestinal villi at 24-48 h post-infection in the human-relevant InlA/B permissive gerbil model

A previous study used a ligated ileal loop model in transgenic-E-cadherin mice and showed *Lm* translocation within 30-45 min post-infection (13). The ligated loop bypasses the stomach; thus, it does not mimic the natural route of gastrointestinal *Lm* infection or allow a follow-on intestinal infection at a later stage. How rapidly *Lm* can translocate across the intestinal epithelium *in vivo* is unknown. Therefore, we first determined the kinetics of *Lm* translocation across the intestinal barrier in the human-relevant InlA/E-cadherin and InlB/Met permissive gerbil model (20).

Immunostaining was used to assess *Lm* association at the intestinal villi in gerbils orally challenged with a single dose (3×10^8^ CFU) of WT clinical strain (F4244, serovar 4b, CC6) at 6, 12, 24, and 48 h post-infection (hpi). Given the rapid kinetics of *Lm* translocation in a ligated ileal loop model (13,14), to our surprise, no ileal or colonic villi-associated *Lm* was observed until 12 hpi (**Fig. 1A-C** and **S1A-B**). *Lm* cells were trapped at the inner mucus layers and remained approximately 20-25 µm away from the ileal and colonic villi until 12 hpi (**Fig. 1A-C** and **S1A-B**). At 24 hpi, only ∼1 in 100 (1%) villi showed *Lm* association with intestinal epithelial cells (IECs) (**Fig. 1A-C** and **S1A-B**). At 48 hpi, ∼30% of the ileal and colonic villi showed *Lm* translocation across the IECs of the villi into the underlying lamina propria (**Fig. 1A-C** and **S1A-B)**. In contrast, *Lm* translocation across the M-cells of the Peyer’s Patches (PP) was observed as rapidly as 6 hpi, and *Lm* counts continued to increase until 48 hpi (**Fig. S2A)**. However, no M cells-associated *Lm* was found in the intestinal villi at all-time points tested (6-48 hpi) (**Fig.** S2B).

**FIG 1.**
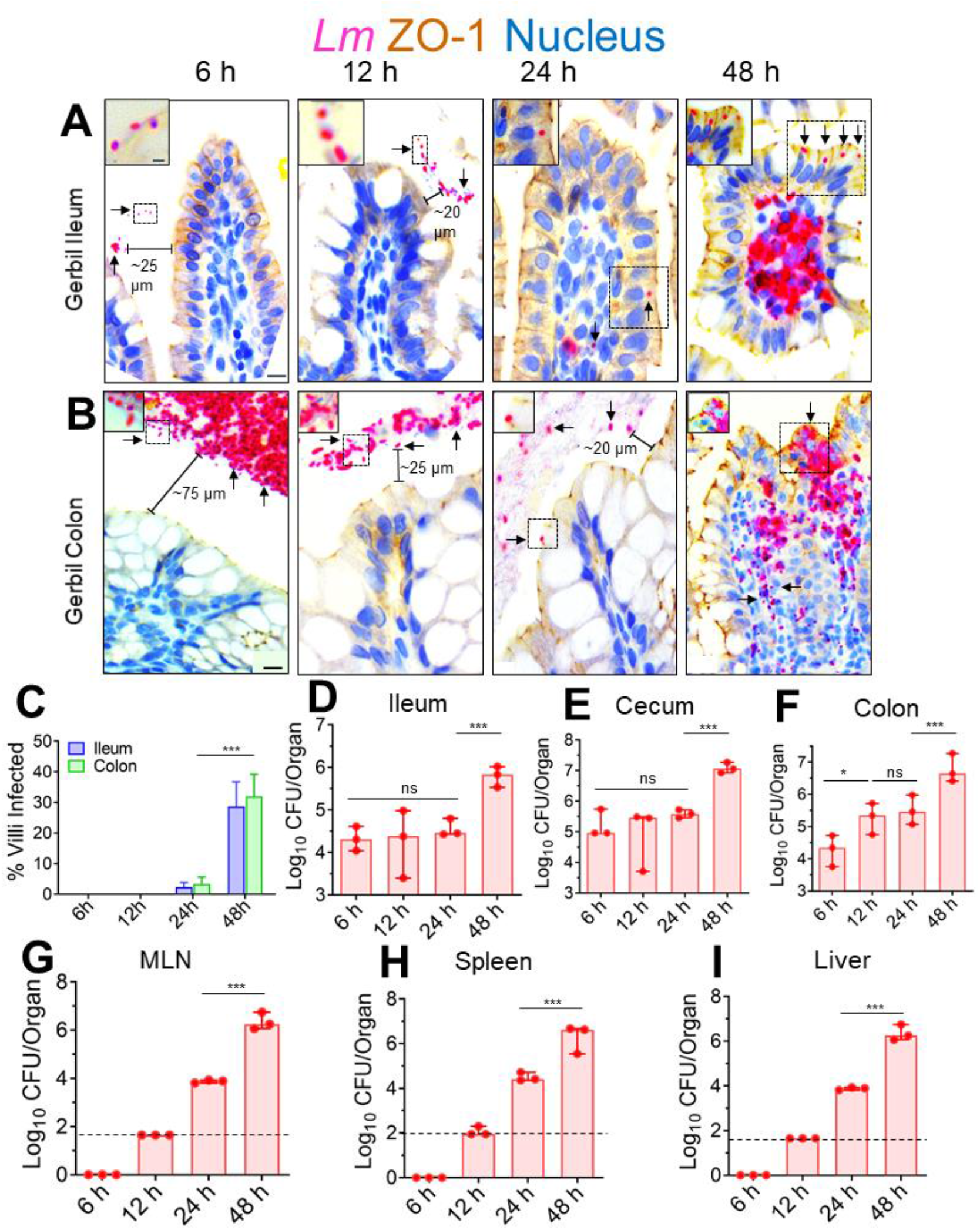
Kinetic analysis of the intestinal invasion of *L. monocytogenes* in orally infected gerbils. **(A-B)** Representative micrographs of ileal (**A**) or colonic villi (**B**) dual immunostained for ZO-1 (tight junction, brown) and *Listeria* (red, arrows) and counterstained with hematoxylin to stain the nucleus (blue) from gerbil orally challenged with ∼3×10^8^ CFU of WT clinical strain (F4244, serovar 4b, CC6) at 6, 12, 24, and 48 hpi; Bars, 10 µm. The boxed areas were enlarged; bars, 1 µm. Translocated *Lm* is observed in the lamina propria (arrows) at 48 hpi but confined in the lumen (arrows) at 6-24 hpi. **(C)** The graph representing quantitative measurements of infected ileal or colonic villi (%) ± SEM. % of infected villi from 100 villi from a single gerbil, 3 gerbil per group, n=300 villi. (**D-I**) *Listeria* counts (total CFU) in the intracellular location in the ileum (**D**) cecum (**E**) and colon (**F**), D-F; gentamicin resistant CFU, and the mesenteric-lymph node (MLN, **G**), spleen (**H**) and liver (**I**) at 6-48 hpi from two-three independent experiments. Dashed horizontal lines indicate the detection limit. Data (**C-I**) represent mean ± SEM of n= 3 gerbils per treatment from three independent experiments. Each point represents a single gerbil. ***p < 0.001; **p < 0.01; *p < 0.05; ns, no significance.

In line with our microscopic observations, at 48 hpi, we observed a significant increase (90-95%) in *Lm* burdens that invaded the ileum, cecum, and colon (**Fig. 1D-F**; gentamicin-resistant colony forming units; CFU). Similarly, bacterial loads increased significantly (90-95%) in the MLN, liver, and spleen at 48 hpi (**Fig. 1G-I**). The detailed kinetic analysis of *Lm’s* intestinal invasion in the gerbil suggests that most *Lm* translocates across the intestinal villi into the underlying lamina propria at 48 hpi. In comparison, the M-cells aid translocation across the PP accounts for early (within 6-24 hpi) bacterial systemic dissemination into the peripheral tissues.

### LAP promotes *Lm* translocation across the intestinal barrier in the InlA-permissive gerbil model

As indicated above, IECs at the villi are critical in *Lm* translocation; therefore, we next determined the contribution of LAP and InlA in *Lm* translocation across the intestinal barrier by assessing bacterial burdens in intestinal and peripheral tissues of orally challenged gerbil at 48 hpi (∼30% of villi infected with WT at 48 hpi; **Fig. 1C**) using several defined genotypes (**Fig. S3A**). The intestinal luminal burdens of WT and *lap^─^* strains are similar (**Fig. S3B)**, indicating that the *lap^─^* does not have a fitness defect for persistence in the gut. However, relative to the WT, the *lap^─^* showed a 100-fold reduced intestinal invasion into the ileum, cecum, and colon (gentamicin-resistant CFU, **Fig. 2A-C**, respectively), which further correlated with ∼100-fold reduced dissemination and colonization in the MLN, liver, and spleen (**Fig. 2D-F**).

**FIG 2.**
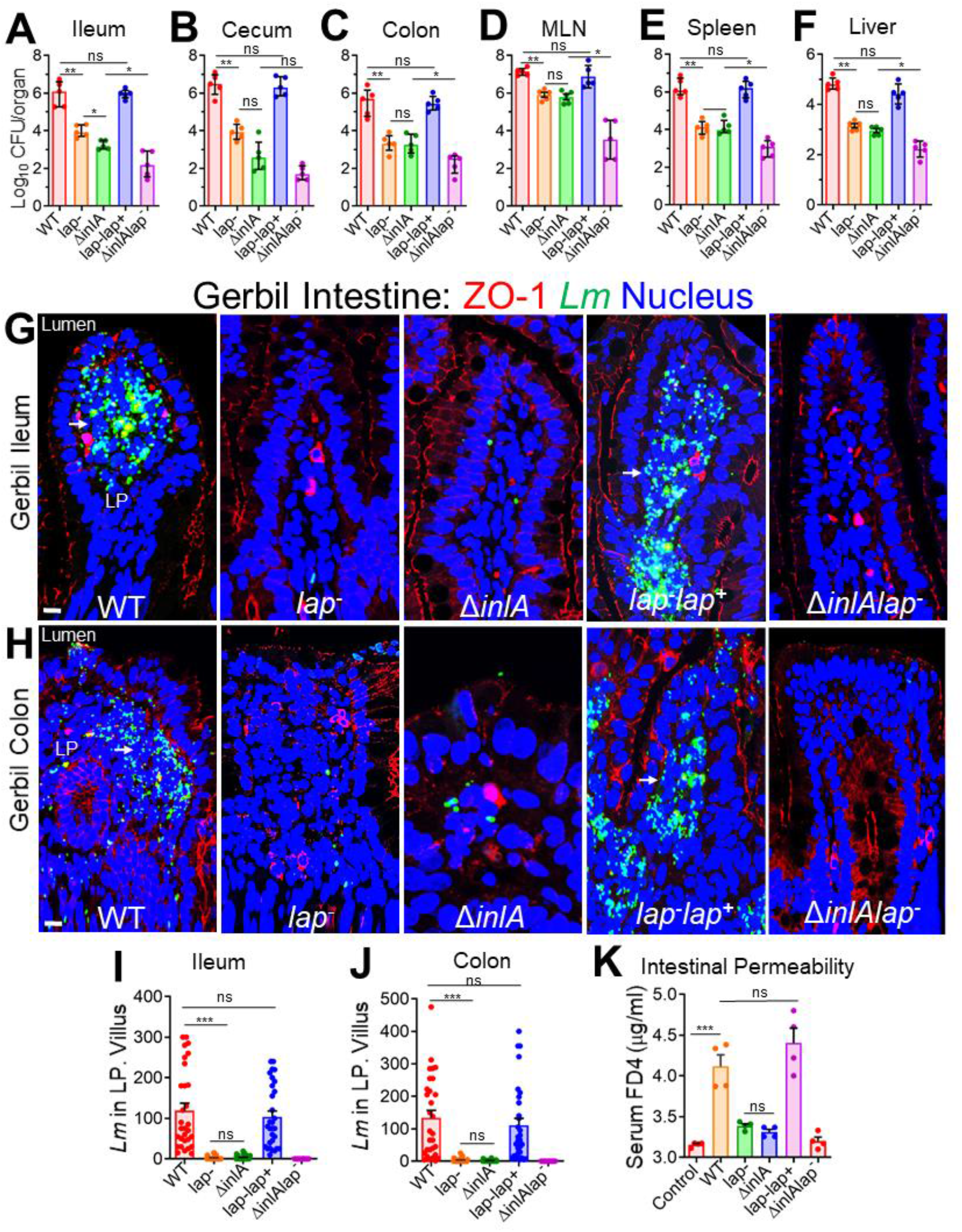
LAP and InlA promote *L. monocytogenes* translocation across the InlA-permissive gerbil intestinal barrier. (**A–F**) Female gerbils were orally challenged with ∼3×10^8^ CFU of WT, *lap*^─^ Δ*inlA*, *lap*^─^ *lap*^+,^ or the Δ*inlA lap*^─^ *L. monocytogenes* bacteria. The plots show the total CFU obtained from the ileum (intracellular) (**A**), cecum (intracellular) (**B**), colon (intracellular) (**C**), mesenteric-lymph node (MLN) (**D**), spleen (**E**), liver (**F**), of gerbils (n=4-6) at 48 hpi from three independent experiments. The bar and brackets represent the mean ± SEM for each group’s data points. All error bars represent mean ± SEM. **p < 0.01; *p < 0.05; ns, no significance. (**G & H**) Representative confocal immunofluorescence microscopic images of ileal (**G**) or colonic (**H**) tissue sections immunostained for ZO-1 (red), *Listeria* (green), and DAPI (blue; nucleus) from WT, *lap*^─^ Δ*inlA*, *lap*^─^ *lap*^+^ or the Δ*inlA lap*^─^ *L. monocytogenes* bacteria-challenged gerbils at 48 h pi, Bars, 10 µm. Increased *L. monocytogenes* (green) was detected in the lamina propria of ileal (**G**) or colonic (**H**) tissue in WT and *lap-lap+*-challenged gerbils (arrows). (**I & J**) Graph representing quantitative measurements of *L. monocytogenes* counts in the lamina propria from ileal (**I**) or colonic (**J**) villi images (n = 30 villi) from three gerbils for each treatment. (**K**) Analysis of 4-kDa FITC-dextran (FD4) permeability through the intestinal epithelium of uninfected (control) and *L. monocytogenes*–infected gerbils in serum at 48 h pi. FD4 was administered orally 4-5 h before sacrifice. Data represent mean ± SEM of n = 4 gerbils per treatment from two independent experiments. All error bars represent mean ± SEM. ***p < 0.001; **p < 0.01; *p < 0.05; ns, no significance.

Immunofluorescence staining of intestinal tissues further confirmed localized translocation defect of the *lap^─^* (∼100-fold reduction) into underlying ileal and colonic lamina propria (**Fig. 2G-J**). Likewise, the *lap^─^* strain also showed 80% less permeability of intestinal barrier paracellular marker, FITC-dextran 4 kDa (FD4), relative to WT gavaged orally to gerbil, 4-5 h before sacrifice (**Fig. 2K**). The *lap^─^lap^+^*complemented strain restored the defects to the WT level (**Fig. 2A-K**).

Consistent with intestinal bacterial burdens, the histopathologic analysis of ileal and colonic tissues revealed increased numbers of polymorphonuclear and mononuclear cells infiltrating the base of the villous lamina propria, crypt cell death, cellular erosion, and increased involvement of the submucosa in gerbils challenged with *Lm* WT relative to *lap^─^* strain while the *lap^─^lap^+^* complemented strain restored the histopathological alterations to the WT levels (**Fig. S3C-F**).

As expected, the Δ*inlA* strain also showed reduced bacterial burdens comparable to the *lap^─^* strain (**Fig. 2A-J**). Notably, a double mutant (*ΔinlAlap^─^*) strain showed fewer bacterial counts than the individual mutant strains (**Fig. 2A-J** and **S3A**), suggesting a detectable additive effect of the two deletions. Similar to WT, both the *lap^─^* and the Δ*inlA* were found in the ileal PP (**Fig. S3G**), suggesting translocation of *Lm* across the PP is LAP and InlA-independent. These data strongly indicate that LAP promotes *Lm* translocation *in vivo* across the intestinal barrier in a human-relevant InlA/B permissive gerbil model.

### LAP targets the classical absorptive enterocytes for *Lm* translocation in the InlA-permissive gerbil model

Next, we determined the relative contribution of LAP and InlA in *Lm* translocation across the three IEC cellular locations: the mucus-secreting goblet cells (GC) in which the mucus is stained by Alcian blue (light blue), the extruding cells (EC) immunostained for cleaved caspase 3 (CC3, brown), and IEC enterocytes (**Fig. 3A-C**). Surprisingly, we found only a very few mucus-secreting goblet cells (**Fig. 3A**, arrows; and **3D**, top panel), and the extruding cells (**Fig. 3B**, arrows; and **3D**, middle panel) were associated with *Lm* WT; the two cell types that are proposed to provide quick InlA access to basolateral E-cadherin (13,14,21). As expected, due to transient apical E-cadherin availability at mucus-secreting goblet and the extruding cells by intrinsic host mechanisms, the association of *Lm* with these IEC subsets was InlA-dependent as the Δ*inlA* showed minimal association with these cells (**Fig. 3A, 3B & 3D,** top and middle panels). The InlA-dependent association of *Lm* with the mucus-secreting goblet and the extruding cells was not due to the changes in their population as similar numbers of these cell subsets were found in the intestine of gerbils challenged with Δ*inlA*, relative to those challenged with WT or other *Lm* strains (**Fig. S4A & B**).

**FIG 3.**
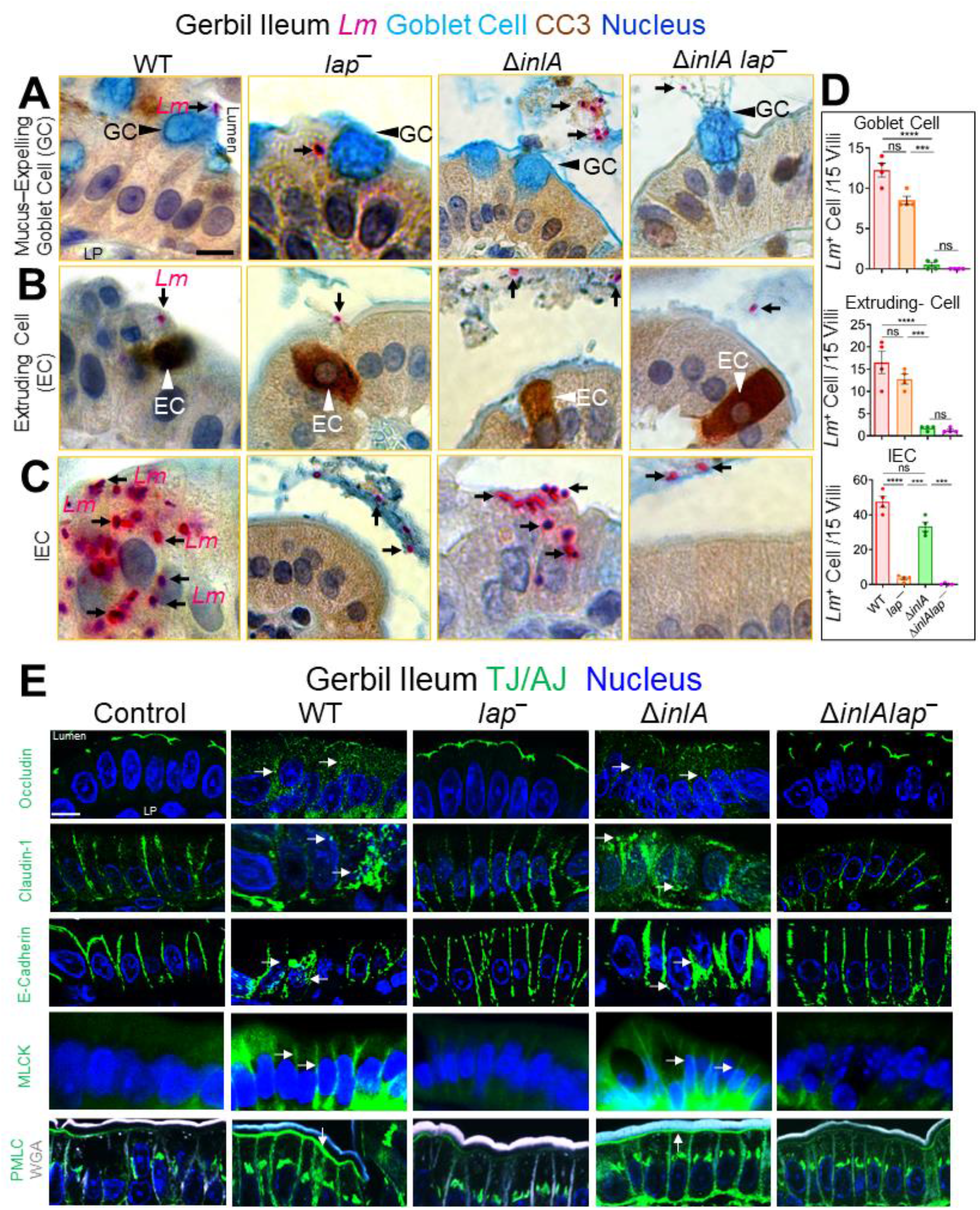
*L. monocytogenes* translocation across absorptive IEC with luminally inaccessible E-cadherin is LAP-dependent. (**A-C**) Representative picto-micrographs of ileal tissue sections dual immunostained for *Lm* (pink) and cleaved-caspase-3 (brown; extruding cells), and Alcian-blue stained for delineating goblet-cell (blue) from *L.monocytogenes* WT, *lap^─^* or Δ*inlA*–infected gerbils at 48 hpi. *Lm* (black arrows) association at and translocation across goblet cell (A, black arrowhead; GC) and extruding cells in different phages of extrusion (B, white arrowhead; EC) is InlA-dependent as *ΔinlA* stain shows no association at these sites. *Lm* (black arrows) association at and translocation across IEC’s (C) with luminally inaccessible E-cadherin is LAP dependent as *lap*^─^ strain shows no association at these sites. The *ΔinlAlap^─^* strain shows negligible *Lm* association at all the cellular sites. (**D**) Graph representing quantitative measurements of *L. monocytogenes* infected cells (in each cell type or at each site) of villi images (n = 60 villi). Each point represents a single gerbil. All error bars represent SEM. ****p < 0.0001; ***p < 0.001; ns, no significance. (**E**) Confocal immunofluorescence micrographs of the ileal tissue sections showing mislocalization (intracellular puncta, endocytosis) of claudin-1, occludin, and E-cadherin (green; arrows), and increased expression of MLCK and P-MLC (green; arrows) in WT, and ΔinlA-infected gerbils but intact localization of occludin, claudin-1 and E-cadherin and baseline expression of MLCK and P-MLC and in *lap^─^,* or the *ΔinlAlap^─^* challenged gerbils (arrows). Images are representative of five different fields from n=3-4 gerbil per treatment. Scale bars, 10 μm. LP, Lamina Propria.

Notably, *Lm* association and translocation were mainly observed at classical IEC enterocytes (**Fig. 3C**, arrows, and **3D**). Furthermore, *Lm* translocation across these enterocytes was LAP-dependent as the *lap^─^* but not the Δ*inlA* strain showed a ∼50-fold reduction in *Lm* attachment and translocation relative to WT (**Fig. 3C,D** bottom panel; arrows). As expected, the double *ΔinlAlap^─^* mutant strain showed negligible bacterial association with either cell type, i.e., mucus-secreting goblet cells, extruding cells, and IEC enterocytes, and the *ΔinlAlap^─^* were observed restricted to the intestinal lumen (**Fig. 3A-D**; arrows). Immunostaining of intestinal tissues for villin, an enterocyte-specific marker, confirmed the classical absorptive enterocyte subtype of *Lm*-associated IECs (**Fig. S4C**). Moreover, the number of *Lm* cells associated with these infected IEC enterocytes was significantly higher (50% increase) than those associated with mucus-secreting goblet or extruding cells (**Fig. S4D**). In another approach, a similar LAP-dependent *Lm* translocation was observed across IEC enterocytes that were stained with (i) wheat germ agglutinin (WGA-white to delineate mucus-secreting goblet cells), which binds to the mucus and carbohydrate residues on the plasma membrane of classical enterocytes at the microvilli, and (ii) 4′,6-diamidino-2-phenylindole (DAPI) to delineate the nuclei of extruding cells at the villi tip (**Fig. S4E**). Predictably, we also found *Lm* cells in the intestinal lumen adjacent to extruded cells and mucus expelled from goblet cells, suggesting the role of epithelial cell shedding and mucus exocytosis in *Lm* clearance (**Fig. S4F**).

Consistent with LAP-dependent *Lm* translocation across absorptive IEC enterocytes, we observed severe mislocalization (endocytosis) of junctional proteins in the ilea of WT and Δ*inlA*-infected gerbils at 48 hpi as revealed by a significantly increased number of cells containing intracellular puncta of occludin (3-fold), claudin-1 (4-fold), and E-cadherin (2-fold) (**Fig. 3E**, arrows; **Fig. S4G**) and significantly increased expression of MLCK (3-fold) and P-MLC (4-fold) (**Fig. 3E**, arrows; **Fig. S4H**). By contrast, ilea of the *lap^─^* and *ΔinlAlap^─^* showed firm localization of the cell-cell junctional proteins and basal levels of MLC phosphorylation, similar to uninfected controls.

In contrast to the active *Lm*-LAP-dependent remodeling of apical junctions at IEC enterocytes (**Fig. 3E**), a relative redistribution of apical TJ protein occludin was observed in the lateral membranes at mucus-secreting (Muc2^+^) goblet cells (**Fig. S5A**) and the extruding cells (a “V” shape TJ) at the tip of the villi (**Fig. S5B**) irrespective of *Lm* infection implying intrinsic host mechanisms. The lateral redistribution of apical occludin was consistent with apical enrichment and availability of E-cadherin during mucus exocytosis (**Fig. S6A**) and apoptotic cell shedding at the tip of the intestinal villi (**Fig. S6B**). These data demonstrate that LAP promotes *Lm* translocation across IEC’s enterocytes that do not usually express luminally accessible E-cadherin in the InlA-permissive host by remodeling and endocytosis of the cell-cell junctional complex.

### Inhibition of caveolin in cells prevents LAP-induced junctional endocytosis, intestinal permeability, barrier loss, and *Lm* translocation

The transport of TJs & AJs to and from the plasma membrane has been reported to occur via four endocytic pathways: macropinocytosis, clathrin-coated, caveolin-mediated, and dynamin-dependent (7). Therefore, we next aimed to understand the mechanism of LAP-induced junctional endocytosis and consequent intestinal permeability, leading to *Lm* translocation. We first used several structurally unrelated inhibitors for each target in Caco-2 cells, as these are unlikely to have overlapping off-target effects. The macropinocytosis inhibitor amiloride and chlorpromazine (CPZ) and pitstop-2, which inhibit endocytosis via clathrin-coated pits, had minimal impact on *Lm* WT-mediated drop in Caco-2 TEER, increase in FD4 flux and *Lm* invasion and translocation (**Fig. 4A-D**). In contrast, methyl-β-cyclodextrin (MβCD) that inhibits caveolae-like membrane domains significantly blocked *Lm* WT-mediated decrease in Caco-2 TEER (∼95%), increase in FD4 flux (∼95%) and *Lm* invasion (∼50%) and translocation across Caco-2 (∼90%) (**Fig. 4A-D**). In addition, the synthetic glycosphingolipid L-t-LacCer (β-d-lactosyl-N-octanoyl-l-threo-sphingosine) that blocks caveolar endocytosis without some of the off-target effects of MβCD (37) showed similar *Lm* blocking (**Fig. 4A-C**). Both clathrin-coated pits and caveolae are required for dynamin-dependent endocytosis. Dynasore inhibiting dynamin (38) also prevented *Lm* WT-mediated decrease in Caco-2 TEER, increased FD4 flux, *Lm* invasion, and translocation (**Fig. 4A-D**). The MβCD, L-t-LacCer, and dynasore restored the purified LAP-mediated (**Fig. S7A**) drop in Caco-2 TEER and elevated FD4 flux (**Fig. 4E, 4F**). Moreover, MβCD, L-t-LacCer, and dynasore also prevented *Lm* WT-induced occludin endocytosis (**Fig. 4G**). These inhibitors independently did not affect Caco-2 TEER (**Fig. S7B**). These data indicate that LAP-induced TJ/AJ endocytosis requires dynamin and involves membrane lipid domains, consistent with a role for caveolae-like membrane domains in this process.

**FIG 4.**
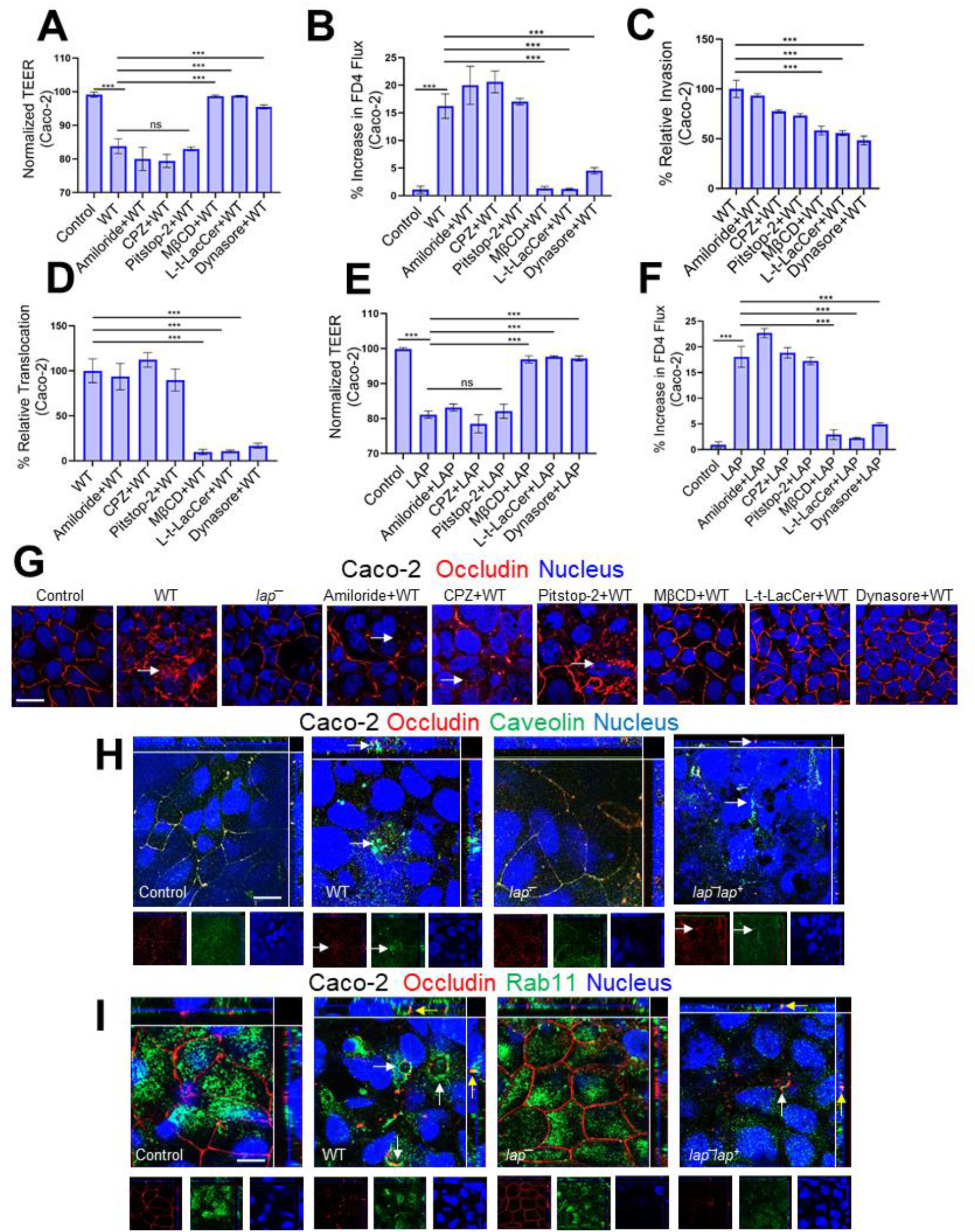
Inhibition of caveolin in cells prevents LAP-induced junctional endocytosis, barrier permeability, and *Lm* translocation. (**A & B**) Trans-epithelial electrical resistance (TEER) measurement of Caco-2 cell monolayer in transwell filter-insert treated pretreated with endocytic pathway inhibitors before *Lm* exposure (MOI; 50, 2h) (**A**) and on the apical (AP)-to-basolateral (BL) flux of paracellular marker 4-kDa FITC-dextran (FD4) permeability (**B**). (**C-D**) Decreased *Lm* invasion (**C**) and translocation (**D**) at MOI of 50 through polarized Caco-2 cell monolayers following pre-treatment with MβCD (10 µM, 30 min), Lt-Lacer (10 µM, 30 min), Dynasore (10 µM, 30 min). (**E & F**) Trans-epithelial electrical resistance (TEER) measurement of Caco-2 cell monolayer in transwell filter-insert treated pretreated with endocytic pathway inhibitors before purified LAP exposure (2 µg/ml, 2 h) (**E**) and on the apical (AP)-to-basolateral (BL) flux of paracellular marker 4-kDa FITC-dextran (FD4) permeability (**F**). (**G**) Representative confocal immunofluorescence micrographs showing intact localization of occludin in Caco-2 cells pretreated with MβCD (10 µM, 30 min), Lt-Lacer (10 µM, 30 min), Dynasore (10 µM, 30 min) before *Lm* exposure (MOI; 50, 45 min). Arrows depict the internalization of occludin. (**H-I**) Representative confocal immunofluorescence micrographs showing colocalization of internalized occludin in Caco-2 following *Lm* WT and *lap^─^lap^+^* but not *lap^─^* exposure (MOI; 50, 45 min) with Caveolin (H, arrows) and Rab11 (I, arrows). Separated channels are shown individually at the bottom of the merged images for clarity.

We next examined markers of different endocytic routes using double immunofluorescence staining. After 45 min post-infection, we observed significant colocalization of internalized occludin (TJ marker) with caveolin-1 and dramatic reorganization of caveolin-1 in Caco-2 cells infected with WT *Lm* and *lap*^─^*lap^+^* but not with *lap*^─^ (**Fig. 4H & S7C**). Significant colocalization of LAP-mediated internalized occludin was also observed with Rab11 (**Fig. 4I & S7D**) and early endosomal antigen 1 (EEA-1) (**Fig. S7E**), the recycling and early endosomal markers. These results suggest that *Lm* LAP-mediated internalized occludin is delivered in early/recycling endosomal compartments.

### Caveolin-1 is required for LAP-induced *Lm* translocation, junctional remodeling, and barrier dysfunction in vivo

Genetic tools in gerbils are limited. Therefore, we used a mouse model to evaluate the role of caveolin-1 *in vivo* in LAP*-*induced junctional remodeling and bacterial translocation. To circumvent the animal species barrier, we created a *lap^─^* mutant in a murinized InlA (InlA^m^) WT, which binds murine E-cadherin with similar affinity as InlA-E-cadherin interaction in humans (**Fig. S8A**) (39,40). Next, we orally challenged caveolin-1 knock-out mice (Cav-1^-/-^) or MLCK knock-out mice lacking the 210-kDa long-chain (MLCK^-/-^) and its parental strain (C57BL/6 WT mice; Cav-1^+/+^ MLCK^+/+^) with InlA^m^, InlA^m^ *lap^─^* or the Δ*inlA* strains and enumerated bacterial burdens in tissues. The total number of InlA^m^, InlA^m^*lap^─^* or the Δ*inlA* strains present in the intestinal luminal contents of the WT mice, the Cav-1^-/-^ and the MLCK^-/-^ mice were similar, indicating that the InlA^m^ *lap^─^*does not possess any fitness defect for survival in the intestinal tract (**Fig. 5A**). However, the WT mice challenged with the InlA^m^*lap^─^* or the Cav-1^-/-^ and the MLCK^-/-^ mice challenged with either the InlA^m^ or the InlA^m^*lap^─^* strain showed a 10-100-fold defect in intestinal invasion into the ileum, cecum, and colon (gentamicin-resistant CFU, **Fig. 5B-D**), relative to WT mice that were challenged with InlA^m^ or the Δ*inlA* strain. Immunostaining of intestinal tissues confirmed ∼200-fold defect in translocation of the InlA^m^*lap^─^*strain in WT mice and the InlA^m^ or the InlA^m^*lap^─^* strain in the Cav-1^-/-^ and the MLCK^-/-^ into the underlying lamina propria, relative to WT mice that were challenged with InlA^m^ or the Δ*inlA* strain (**Fig. 5E, 5F, S8B & S8C**).

**FIG 5.**
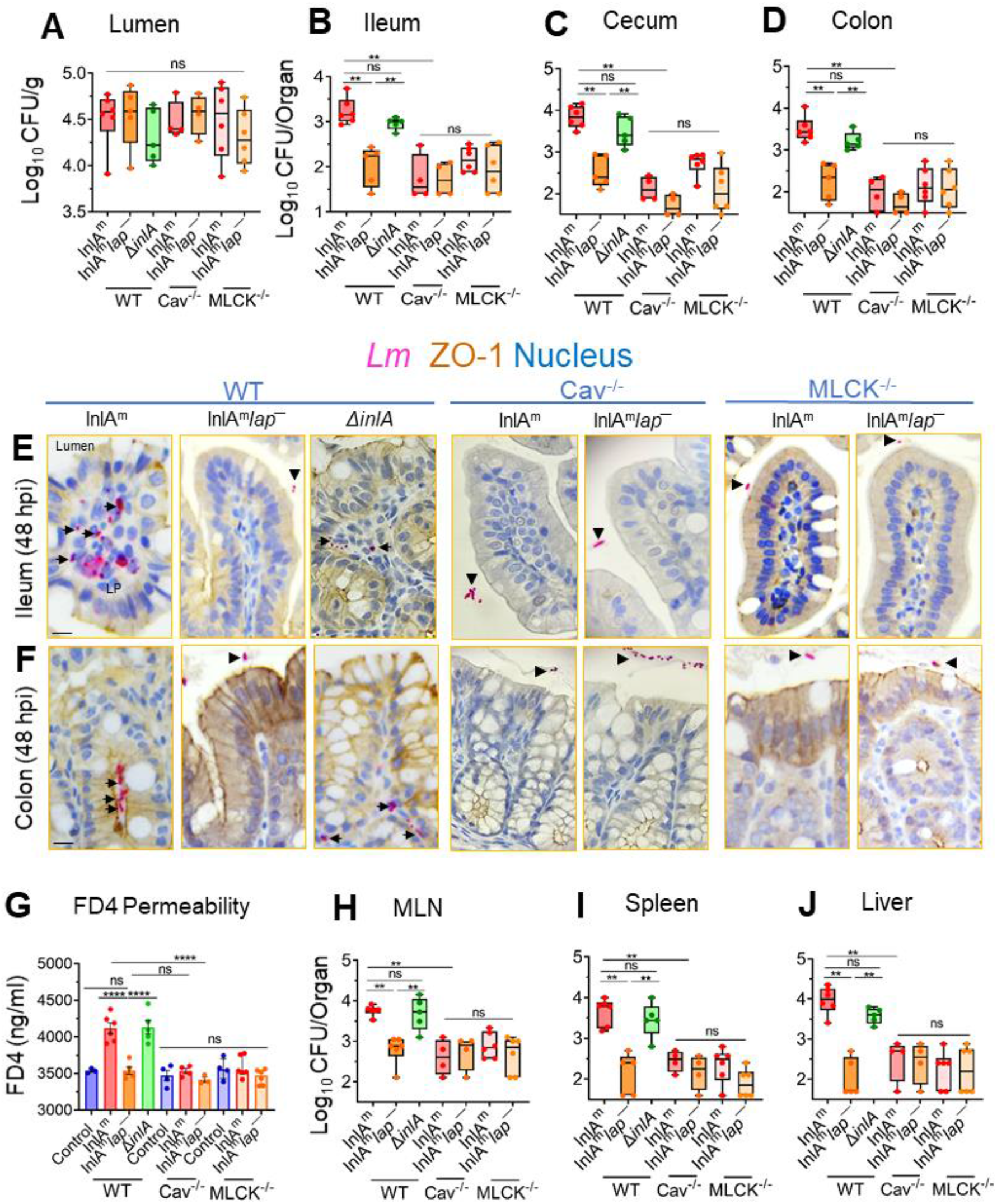
*Listeria monocytogenes* translocation and intestinal epithelial permeability are affected in caveolin-1 knockout mice. (**A-D**) Male and female wild-type C57BL/6 (Cav^+/+^, MLCK^+/+^) or the caveolin-1 ^-/-^ knockout mice (Cav^-/-^) or the 210-kDa MLCK knockout mice (MLCK^-/-^) mice were orally gavaged with 5×10^8^ CFU of InlA^m^, InlA^m^*lap*^─^ or Δ*inlALm*. The box plot shows the total CFU obtained from the lumen (A) ileum (intracellular) (B), cecum (intracellular) (C), colon (intracellular) (D) of mice (n=4-5) at 48 hpi from three independent experiments. The bar and brackets represent the median ± range for each group’s data points. Bar and brackets represent the median ± range. (**E & F**) Representative microscopic images of ileal (**E**) or colonic (**F**) tissue sections immunostained for ZO-1 (brown) *Listeria* (pink) from *Lm*-challenged mice at 48 h pi, Bars, 10 µm. Arrows denote increased *Lm* detected in the lamina propria of ileal or colonic tissue in wild-type C57BL/6 mice challenged with InlA^m^ or *ΔinlA*. Arrowheads denote bacteria restricted to the intestinal lumen. (**G**) Analysis of paracellular permeability of 4-kDa FITC-dextran (FD4) through the intestinal epithelium of uninfected (cont) or *Lm*–infected, WT (MLCK^+/+^ Cav^+/+^), the Cav ^-/-^ or the MLCK-/- mice in serum at 48 h pi. FD4 was administered 4-5 h before sacrifice. Data represent mean ± SEM of 3-6 mice per treatment from three independent experiments. (**H-J**) The box plots show the total CFU obtained from the mesenteric-lymph node (MLN) (H), spleen (I), liver (J) (n=4-6) at 48 hpi of *Lm*–infected, WT (MLCK^+/+^ Cav^+/+^), the Cav ^-/-^ or the MLCK^-/-^ mice. Bar and brackets represent the median ± range, respectively, for the data points in each group. ***, P<0.001; **, P<0.01; *, P<0.5; ns, no significance.

To assess whether decreased bacterial translocation correlates with decreased paracellular permeability, we examined FD4 permeability through the intestinal epithelium. Relative to uninfected control WT mice, *Lm*-infected mice that received FD4 orally 4-5 before sacrifice displayed significantly increased FD4 concentrations (∼15 %) in the serum of WT mice challenged with the InlA^m^ or the Δ*inlA* strain (**Fig. 5G**). However, the FD4 concentrations were significantly lower in the serum of WT mice challenged with the InlA^m^ *lap^─^* or the Cav-1^-/-^ and the MLCK^-/-^ mice challenged with InlA^m^ or the InlA^m^ *lap^─^* and were similar to the uninfected controls. The impaired intestinal translocation and permeability of the InlA^m^ *lap^─^*strain in WT mice or the InlA^m^ and InlA^m^ *lap^─^* strain in the Cav-1^-/-^ and the MLCK^-/-^ mice correlated with significantly reduced (∼100-fold) bacterial dissemination to the MLN, liver, and spleen relative to WT mice that were challenged with InlA^m^ or the Δ*inlA* strain (**Fig. 5H-J**).

Consistent with LAP, Cav-1, and MLCK-dependent bacterial translocation and intestinal permeability, immunofluorescence staining of the intestinal tissue sections revealed membrane mislocalization (endocytosis) of claudin-1 (3-4 fold), occludin (3-fold), and E-cadherin (2-fold) and increased expression of P-MLC (3-fold) in WT mice challenged with the InlA^m^ or the Δ*inlA* strain (**Fig. S8D & S8E**). In contrast, we observed firm localization of these cell-cell junctional proteins in the WT mice challenged with the InlA^m^ *lap^─^* strain or the Cav^-/-^ and the MLCK^-/-^ mice challenged with either the InlA^m^ or the InlA^m^ *lap^─^*. Similar increases in epithelial MLC phosphorylation (P-MLC) were observed in WT and Cav^-/-^ mice challenged with the InlA^m^ strain, suggesting MLCK activation precedes caveolin-1-mediated junctional endocytosis (**Fig. S8D & S8F**). These data demonstrate that LAP-mediated *Lm* translocation requires the intestinal epithelial caveolin–1–dependent junctional endocytosis.

### LAP-mediated endocytosis of epithelial junctions provides easier InlA access to the basolateral E-cadherin for *Lm* translocation

We next determined whether LAP directly facilitates InlA-mediated invasion by providing *Lm* InlA access to basolateral E-cadherin. We coated *L. innocua* (*Lin,* nonpathogen) with exogenously purified InlA (Lin^InlA^, **Fig. S9A**) and coinfected the human polarized Caco-2 or HCT-8 cells with a 1:1 ratio of *Lm*WT ^(emR)^: Lin^InlA^ ^(emS)^ or Lm *lap^─^* ^(emR)^: *Lin*^InlA(emS)^ or (**Fig. 6A,B)**. Relative to *Lm lap^─^,* LAP in the WT during coinfection promoted 90% of InlA-mediated invasion of the Lin^InlA^. Consistent with these data, the invasion into and translocation across Caco-2 and HCT-8 of the *lap^─^* showed a 50 and 100-fold defect (**Fig. 6C,D)** relative to the WT strain. As expected, the invasion and translocation of the *ΔinlAlap^─^* double mutant strain was severely attenuated.

**FIG 6.**
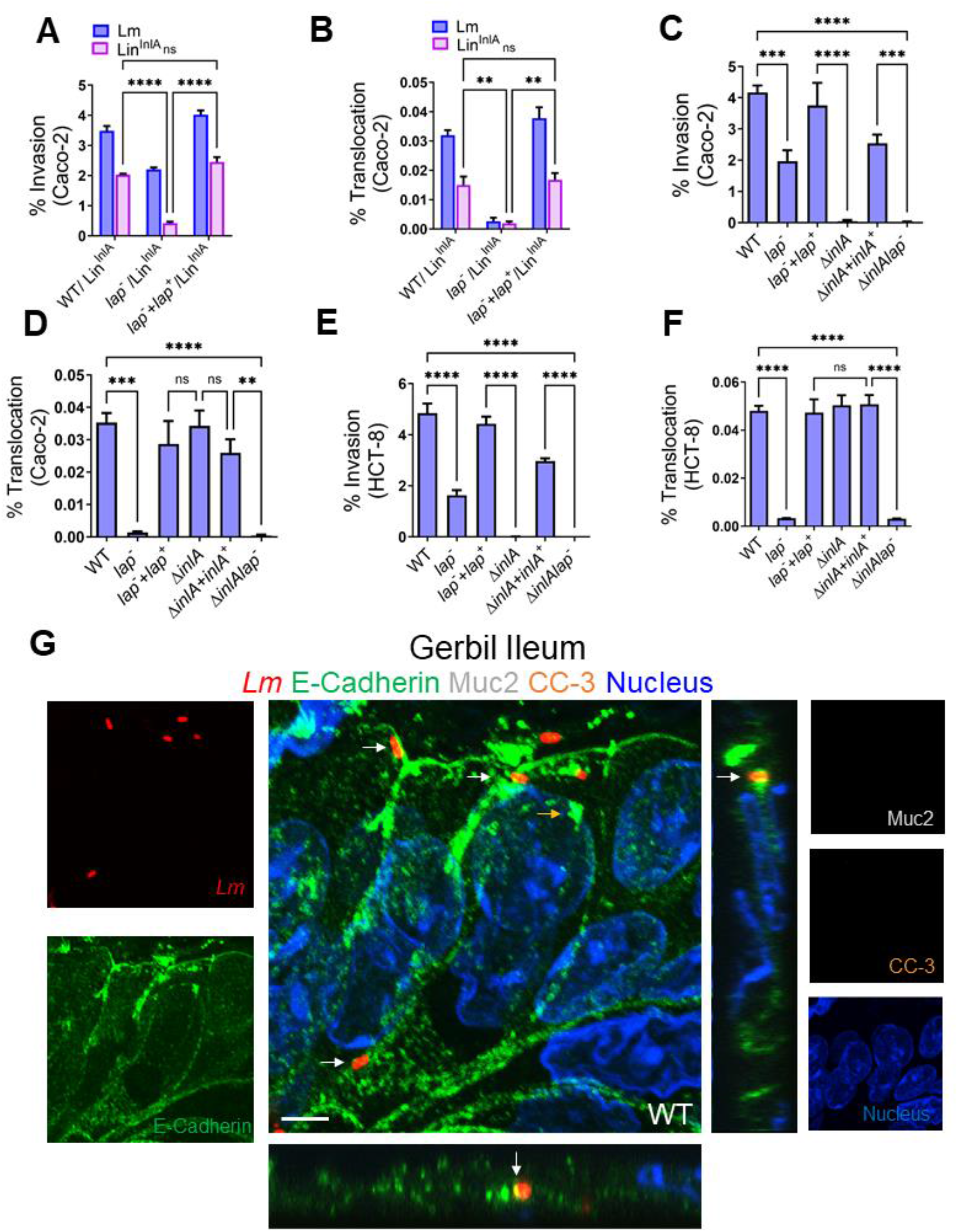
LAP-mediated endocytosis of epithelial junctions provides easier InlA access to the basolateral E-cadherin for *Lm* translocation. (**A & B**) Analysis of Lin^InlA^ invasion of Caco-2 and HCT-8 cells co-infected with a 1∶1 (MOI; 50) mixture of Lin^InlA^ and *Lm* WT or Lin^InlA^ and *lap*^─^. (**C-F**) Analysis of invasion (**C&D**) and translocation (Transwell filter-inserts) (**E&F**) of *Lm* WT and isogenic strains in polarized Caco-2 and HCT-8 cell monolayers infected at an MOI of 50. Data represent mean ± SEM from three independent experiments, n=6. (**G & H**) Analysis of Δ*inlAlap*^─^ invasion in Caco-2 (**G**) and HCT-8 (**H**) with exogenously added LAP (Δ*inlAlap*^─^+LAP), InlA (Δ*inlAlap*^─^+InlA), or Δ*inlAlap*^─^+LAP & InlA. **(I)** Representative confocal immunofluorescence picto-micrographs of the gerbil ileal tissue sections immunostained for E-cadherin (green), *Listeria* (red; arrows), Muc-2 (White, Goblet Cell), Cleaved Caspase-3 (orange), and DAPI (blue; nucleus) from WT challenged gerbil at 48 h pi. Bars, 5 µm. *Lm* (arrows, yellow in merged images) was observed attached to the IECs (Muc2 and CC3) and exiting into lamina propria in WT with co-localized E-cadherin at the adherens junctions. Separated channels are shown individually to the left or right) of the merged images. The X-Z and Y-Z cross-sections were produced by orthogonal reconstructions from the z-stack scanning. Pictures are representative of five different fields from three gerbils. LP, Lamina Propria. ***, *P*<0.001; **, *P*<0.01; *, *P*<0.5; ns, no significance.

Immunostaining of gerbil intestinal tissues for *Lm,* Muc-2 (to delineate the mucus-secreting goblet cells), cleaved caspase-3 (CC3) (to delineate extruding cells) further confirmed colocalization of *Lm* with E-cadherin and active *Lm* translocation across IEC’s in the absence of the mucus-secreting goblet cells and extruding cells (**Fig. 6B,G**). In contrast, the *lap^─^*mutant strain showed a negligible association and a significant defect in translocation across these IEC’s (**Fig. 4E**). InlA access to E-cadherin at the cell-cell adherens junctions of enterocytes is also evident in the gerbil intestinal tissues immunostained for *Lm* and E-cadherin (**Fig. S9B**). These data demonstrate that the LAP provides easier access of InlA to E-cadherin at the enterocyte-cell-cell-cell junction for InlA-mediated *Lm* epithelial invasion by endocytosis of apical junctional proteins.

## DISCUSSION

Breach of host barriers by pathogens is a crucial aspect of the invasive infection process (1–3). Gaining insights into how these microbes breach the intestinal barrier is critical in deciphering the underlying pathogenic mechanisms of foodborne pathogens. *Lm* is a model intracellular pathogen that must initially breach the critical intestinal barrier to gain access for systemic dissemination. Previous work has demonstrated that *Lm* uses the M-cell in Peyer’s patch (41) and the InlA-mediated pathways (13,14) to invade and cross the intestinal barrier. Paradoxically, *Lm* crosses the intestinal barrier in animal models where both paths are absent, such as in Peyer’s patch-null mice (17,41) or defective E-cadherin in mice (16).

Additionally, *Lm* isolates from human clinical cases have been found expressing a defective InlA (truncated protein) (24,25) and yet infect human newborns and fetuses of pregnant guinea pigs after oral dosing (26,27). These observations strongly suggest InlA- and M-cell-independent routes for *Lm* to cross the intestinal barrier. The InlA-independent *Lm* invasion mechanism is understudied and significant for a comprehensive understanding of the *Lm* translocation process.

We previously reported an InlA-independent translocation of *Lm,* which requires the engagement of LAP with its apically expressed receptor Hsp60, leading to the activation of canonical NF-κB(p65) signaling, thereby promoting the MLCK-mediated opening of the intestinal cell-cell junction for bacterial translocation into the lamina propria and systemic dissemination in a mouse model (6). Mice are non-permissive to InlA-E cadherin interactions (16); therefore, the precise contribution of LAP *in vivo* in an InlA-permissive host (as humans) is unknown. Here, we studied the relative contribution of LAP and InlA in *Lm* crossing of the intestinal barrier *in vivo* in a Mongolian gerbil model that is permissive to both InlA/E-cadherin and InlB/Met systems (20), which allowed us to study the pathophysiology of disease in animal models relevant to human listeriosis.

Previous work has identified that the interaction of InlA with accessible E-cadherin sites allows rapid (within 30–45 min) InlA-dependent translocation of *Lm* in the ligated jejunal loops in transgenic mice expressing human E-cadherin (13). Notably, the ligated loop model is distinct from the natural gastrointestinal route of *Lm* infection, bypassing the stomach. Consequently, it does not accurately replicate the typical path and kinetics of gastrointestinal *Lm* infection.

Following oral inoculation, our results in the InlA-permissive gerbil model of gastrointestinal infection indicate initial rapid seeding of *Lm* at the M-cells in the Peyer’s patch within 6-12 hpi (**Fig. 1 & S1**). However, *Lm* association and translocation across the intestinal ileal and colonic villi were only discernible at 24-48 hpi. Previous studies reported no significant difference in intragastrically inoculated *ΔinlA* mutant in WT mice and transgenic mice engineered to express ‘humanized’ E-cadherin until 24-48 hpi (20). Furthermore, when mice were challenged with InlA^m^ with high affinity for mouse E-cadherin, there were no significant differences in intestinal *Lm* burdens compared to mice inoculated with *Lm* WT strain until 24-48 hpi (40). No discernible distinctions in bacterial translocation of *ΔinlA* mutant, relative to WT until 24-48 hpi in *Lm* infection models with successful InlA-E cadherin interaction suggesting that the initial rapid translocation of *Lm* is mediated non-specifically by the M-cells in Peyer’s patch and are consistent with our findings.

Our results indicate that both the *lap^─^* and the Δ*inlA* strains were significantly impaired in their ability to cross the intestinal barrier, exit into the underlying lamina propria, and disseminate systemically in gerbil tissues (**Fig. 2**), suggesting that both LAP-dependent paracellular translocation and InlA-mediated transcytosis are required for efficient *Lm* crossing of the intestinal barrier. A double mutant strain (Δ*inlAlap*^─^) exhibited lower bacterial burdens in intestinal and peripheral tissues than the individual mutant strains, indicating a noticeable cumulative effect resulting from both deletions. We further identified that *Lm* uses LAP to translocate across absorptive IECs enterocytes that do not express luminally accessible E-cadherin by directly remodeling the AJC to gain access to the underlying lamina propria for successful infection (**Fig. 3**). Furthermore, in contrast to *Lm* transcytosis facilitated by the InlA-E-cadherin interaction, occurring passively at the extruding cells and empty goblet cells, which are inherent innate defenses that *Lm* utilizes to infiltrate the lamina propria. The LAP-dependent remodeling of enterocyte junctional complex for translocation strongly implies *Lm* employs an active mechanism. Moreover, enterocytes constitute approximately 90% of the cellular composition within the villi (42,43). Therefore, this abundance of enterocytes may afford *Lm-* enhanced prospects for translocation across the intestinal villi, particularly in contrast to the lesser prevalence of goblet cells (5-10%) and extruding cells (1-5%) localized at the distal tips of the villi (42,43).

Pharmacological inhibition studies suggest only dynasore, MβCD, and L-t-LacCer were effective in inhibiting the *Lm* translocation and junctional endocytosis, indicating LAP-mediated junctional endocytosis relies on dynamin and membrane regions rich in cholesterol, such as caveolae (**Fig. 4**). In line with these observations, we found mice lacking caveolin-1 when challenged with InlA^m^ with high-affinity interaction of modified InlA with mouse E-cadherin prevented LAP-induced junctional endocytosis, intestinal barrier permeability, and *Lm* translocation (**Fig. 5**). Notably, the absence of caveolin-1 did not prevent LAP-mediated MLC phosphorylation, indicating that the activation of MLCK occurs prior to caveolin-1-mediated junctional endocytosis (**Fig. 5**). Furthermore, we found intracellular tight junctions to colocalize with caveolin-1, EEA^+^ early endosomes, and Rab11^+^ recycling endosomes in cells exposed to WT strain but not the *lap^─^* strain, underscoring that LAP-induced junctional proteins internalization occurs through caveolin-1-dependent endocytosis and are trafficked to the early and recycling endosomal intracellular destinations (**Fig. 4**). The requirement of LAP in junctional endocytosis and *Lm* translocation is consistent with the need for host Hsp60 and aligns with our previous observations that WT strain induced junctional protein redistribution (6) and was significantly impaired (90%) in Caco-2 with *hsp60* knocked-down, while the *lap^─^* had no change (6,44). Conversely, overexpression of *hsp60* significantly increased the translocation of WT and *ΔinlA* but not *lap^─^* (6,44). Our data suggest a central role of caveolin-1, which necessitates dynamin and cholesterol-enriched membrane domains and the host’s intracellular trafficking machinery, including vesicle transport systems for the LAP-Hsp60 interaction-mediated enterocyte junctional remodeling and *Lm* translocation across the intestinal barrier.

Our results provide a molecular explanation for the respective and complementary contributions of LAP and InlA in *Lm* translocation across the intestinal barrier during infection *in vivo* in human-relevant InlA-permissive hosts. Our data suggest that *Lm* translocation across the enterocyte cell-cell junction is a two-step process in which LAP-mediated remodeling of the apical junction serves as a first critical precursor event for subsequent InlA-dependent epithelial transcytosis that provides access to basolateral E-cadherin at the enterocyte epithelial cell-cell junction (**Fig. 6**), in addition to host innate defense by “villous cell extrusion” (**Fig. 3**) (21) and mucus expelling goblet cell (**Fig. 3**) (13). Additionally, LAP-Hsp60-mediated translocation acts as an active mechanism for InlA-independent paracellular translocation across the enterocyte cell-cell junction (3,6). Multiple reports indicate that *Lm* isolates, primarily from environmental or food sources – expressing truncated InlA can infect humans (24–27); thus, these results have substantial implications for public health and food safety regulations and management of listeriosis in immunocompromised, pregnant, and aging populations.

## MATERIALS AND METHODS

### Bacterial Strains and Growth Conditions

All bacterial strains used in the study are listed in **Table S1**. Unless otherwise indicated, all bacterial strains were grown at 37°C with shaking for 12-16 h. *L. monocytogenes* F4244 (WT; serovar 4b, CC6), the Δ*inlA* in-frame deletion mutant (AKB301)(44) (AKB301), the *inlA*^m^ knock-in mutant strain (39), and *L. innocua* (*Lin*) F4248 were grown in tryptic soy broth containing 0.6% yeast extract (TSBYE; BD Bioscience). The isogenic *lap*-deficient insertional mutant strain in WT background (*lap^—^*; KB208) (30) and the isogenic *lap*-deficient insertional mutant strain in the *inlA*^m^ knock-in mutant background (*inlA*^m^*lap^—^*)were grown in TSBYE containing erythromycin (Em; 5 µg/ml) at 42°C. The *lap*^—^ stain complemented with the *Lm lap* (*lap^—^lap+*; CKB208) (30) was grown in TSBYE containing erythromycin (5 μg/ ml) and chloramphenicol (5 μg/ml) at 37 °C. The erythromycin-resistant *Lm* F4244 were grown in TSBYE containing erythromycin (Em; 5 µg/ml) at 37°C.

### Gerbil and Mice

Female Mongolian gerbils weighing 51-60 g (∼8-10-week-old) were obtained from Charles River (strain 243). C57BL/6 mice, 6–8-week-old, male or female, the caveolae protein 1 (Cav1) knock-out (*Cav-1^−/−^*; the Jackson Laboratory strain # 007083) or the 210-kDa MLCK^−/−^ mice (6,45) or the wild-type C57BL/6J mice, bred in our facility were used (Table S2). Following arrival, gerbils were housed in a group of two gerbils/cage, and mice housed in five mice/cage were provided mouse chow and water at lib. On the day of the challenge, food and water were removed from the cages 12 h before oral gavage to prevent mechanical blockage of the *Listeria* inoculum by food in the stomach, which may cause the inoculum to aspirate into the lungs. The 12-h grown *Lm* WT, *lap^─^*, *ΔinlA*, *lap^─^lap^+^ ΔinlAlap^─^*, InlA^m,^ and InlA^m^ *lap^─^* strains, each resuspended in 200 μl of phosphate-buffered saline (PBS, pH 7.4) containing approximately ∼3×10^9^ CFU for Gerbil and ∼ 5×10^8^ for WT C57BL/6, MLCK^−/−^ or Cav-1^−/−^ mice were administered orally to randomly selected gerbil using a stainless-steel ball-end feeding needle (Popper). The control gerbil or mice received only PBS. The food was returned 1 hpi, and the gerbil or mice were sacrificed 12-48 hpi using CO_2_ asphyxiation. All animal procedure (IACUC Protocol no. 1201000595) was approved by the Purdue University Animal Care and Use Committee, which adheres to the recommendations of the Guide for the Care and Use of Laboratory Animals published by the National Institutes of Health.

### Mammalian cells

The human colon carcinoma Caco-2 cell line (ATCC # HTB37) and the HCT-8 (ATCC # CCL-244) human ileocecal cell line from 25 to 35 passages were cultured in Dulbecco’s Modified Eagle’s medium (DMEM) (Thermo Fisher Scientific) supplemented with 4mM L-glutamine, 1 mM sodium pyruvate, and 10% fetal bovine serum (FBS; Atlanta Biologicals).

### Construction of *Listeria* mutant strains

To generate a *lap* insertion mutant, integrative pPAad101 was constructed by ligating a partial lap of 1 kbp into pAUL-A, a temperature-sensitive shuttle vector (30). To create *ΔinlAlap^─^* or *InlA^m^lap^─^* double mutant, ΔinlA or *InlA^m^*competent cells were transformed with pPAad101, the same plasmid used to generate *lap−* insertion mutant. The transformants were incubated at 30°C overnight to optimize the plasmid integration into the host chromosome. Positive clones were selected from TSBYE agar containing 5 μg/mL erythromycin at 42°C. Protein expression was validated by Western blotting.

### Enumeration of *L. monocytogenes* in Gerbil and Mouse Organs

*Listeria* in the extra-intestinal sites was enumerated in the organs that were harvested aseptically following oral infection and homogenized using a tissue homogenizer in 4.5 ml (spleen and MLN) or 9 ml (liver) of buffered-Listeria enrichment broth (BLEB; Neogen) containing 0.1% Tween 20 and selective antimicrobial agents (Neogen). The samples were serially diluted in PBS and plated onto modified Oxford (MOX; Neogen) agar plates containing selective antimicrobial agents (Neogen). *Listeria* in the intestinal lumen was assessed in the entire intestinal contents, which were removed and homogenized using a tissue homogenizer (Bio Spec) in 5ml of BLEB containing 0.1% Tween 20 (PBS-T). The invaded bacterial counts in the intestinal sites were enumerated in the harvested intestinal segments (ileum, cecum, and colon). Briefly, the segments were rinsed three times in a Perti-dish containing 10 ml Dulbecco’s Modified Eagle’s medium (DMEM) and then incubated for 2 h in 15 ml of DMEM containing 100 µg/ml of gentamycin sulfate to kill the extracellular bacteria in the lumen. The sections were then rinsed three or more times in 15 ml of DMEM and homogenized in a round-bottom tube using a tissue homogenizer in 1 ml BLEB. The samples were serially diluted in PBS and plated onto MOX agar plates supplemented with antimicrobial agents. In specific experiments, small sections of ileal and colonic tissue samples (1 cm) were cut and fixed overnight in 10% formalin for histopathology or immunostaining.

### Immunohistochemistry, Alcian Blue Staining, and Histopathology

Gerbil or Mouse tissues were fixed in 10% neutral buffered formalin for 24–48 h, placed in a Sakura Tissue-Tek VIP6 tissue processor for dehydration through graded ethanol (70%,80%, 95%, and 100%), cleaned in xylene, and embedded in Leica Paraplast Plus paraffin. Tissue sections (4 μm) were made using a Thermo HM355S microtome. Sections were mounted on charged slides and dried for 30–60 min in a 60 °C oven. All slides were deparaffinized through three changes of xylene (5 min each) and rehydrated through graded ethanol as above in a Leica Autostainer XL. Slides are stained in Gill’s II hematoxylin blue and counterstained in an eosin/phloxine B mixture using the Leica Autostainer XL. Finally, slides were dehydrated, cleared in xylene, and mounted with coverslips in a toluene-based mounting media (Leica MM24). For immunohistochemistry, after deparaffinization, antigen retrieval was done in the appropriate buffer using a BioCare decloaking chamber at 95 °C for 20 min. Slides were cooled at room temperature for 20 min and dipped into TBST. The rest of the staining was carried out at room temperature using a BioCare Intellipath stainer. Slides were incubated with 3% hydrogen peroxide in water for 5 min, or Bloxall block for 10 min for antibody labeling. Slides were rinsed with TBST and incubated in 2.5% normal goat or horse serum for 20 min. Excess reagents were removed, and a primary antibody or antibody cocktail was applied at the appropriate dilution for 30 min. Primary antibodies (Table S2)) include antibodies to *Listeria* (1:100 dilution), ZO-1 (1:100 dilution), M-cell (1:100 dilution), cleaved caspase-3 (1:200 dilution), Villin (1:100 dilution), E-cadherin (1:200 dilution). Negative control slides were stained with their respective isotype controls (Table S2) at 1–2 μg/mL for 30 min. After TBST rinse (twice), the secondary antibody was applied for 30 min and rinsed (twice) in TBST before reaction with Vector ImmPACT DAB (Vector Labs) for 5 min. Slides probed with two antibodies were counterstained with ImmPACT Vector Red (Vector Labs). Slides were rinsed in water and counterstained with hematoxylin. Tissue sections were also stained with Alcian blue for goblet cell staining.

Thin-stained hematoxylin and eosin tissue sections from above were examined microscopically by a board-certified veterinary pathologist blinded to the treatment groups, and the interpretations were based on standard histopathological morphologies. The pathologist was blinded to the bacterial strain and compared the ileal or colonic sections to the controls as previously (6).

To determine the extent of the inflammation, the gerbil ileal tissues were scored on a scale of 0-3 for two parameters, yielding a maximum score of 6. The scoring parameters were the amount of polymorphonuclear leukocyte infiltrate and mononuclear infiltrate.

To determine the extent of cellular inflammation and damage, the Gerbil colonic tissues were scored on a scale of 0-3 for five parameters, yielding a maximum score of 15. The scoring parameters were the amount of polymorphonuclear leukocyte infiltrate, mononuclear infiltrate crypt cell death, erosion, and submucosal involvement.

To grade the amount of polymorphonuclear leukocyte infiltrate and mononuclear infiltrate, the following histomorphological scale was used: 3 = markedly increased, 2 = moderately increased, 1 = slightly increased, and 0 = normal. To score the involvement of the submucosa, the following histomorphological scale was used: 3 = 50% or greater of the submucosal diameter, 2 = 10%-50%, 1 = <10%, and 0 = normal.

H&E-stained and immunoperoxidase-stained tissues were imaged using a DMLB microscope (Leica) with ×40/0.25 NA HC FL PLAN or a ×100/1.40 NA HC FL PLAN oil immersion objective and a DFC310 FX (Leica) camera-controlled by Leica Application Suite. Post-acquisition processing, including the stitching of tiled images, was performed using the Leica Application Suite (Leica). Immunoperoxidase-stained positive cells were counted manually on tiled images in a blinded manner. For each experiment, immunoperoxidase-stained positive cells from 15-25 villi in the tissue sections of three to four individual animals per treatment were recorded. Each point represents an average of 15-25 villi from a single mouse/gerbil.

### Immunofluorescence Staining and Confocal Microscopy

The gerbil or mice ileal and colonic tissue sections were collected, fixed with 10% formalin, and embedded in paraffin. The tissues were sectioned (5 µm thick), deparaffinized, and rehydrated for antigen retrieval by immersing the slides in boiling sodium citrate buffer (10 mM, pH 6.0) or 0.01M Tris/EDTA (pH 9.0), for 10 min. The tissue sections were permeabilized and blocked with PBS containing 0.3% Triton X-100 (Sigma-Aldrich) and 5% normal goat serum (Cell Signaling) and immunostained with specific primary antibodies or a cocktail of primary antibodies (Table S2) by incubating overnight at 4°C. Primary antibodies included antibodies to MLCK (1:100 dilution), P-MLC (1:200 dilution), claudin-1 (1:200 dilution), occludin (1:150 dilution), and E-cadherin (1:200 dilution). Tissue sections were incubated with Alexa Fluor-488, Alexa Fluor-555 conjugated secondary antibody with or without Alexa Fluor-647 conjugated-wheat germ agglutinin (WGA; Sigma, which binds to sialic acid and N –acetyl glucosaminyl carbohydrate residues on the plasma membrane of enterocytes and in the mucus of goblet cells for 2 h at room temperature followed by washing three times with PBS (3 cycles, 5 min). The nuclei were stained with DAPI (1 µg/ml; Cell signaling), and slides were mounted in ProLong antifade reagent (Invitrogen) For antibody labeling in cells, Caco-2 cells were grown to 40–50% confluence in eight-chambered slides (Millipore). At the end of the treatment, the cells were fixed with 3.7% formaldehyde in PBS for 20 min and permeabilized and blocked with PBS containing 0.3% Triton X-100 and 3% BSA (Sigma-Aldrich) for 1 h at room temperature and then incubated with primary antibodies to Occludin (1:50 dilution), Caveolin −1 (1:100 dilution), Rab11a, (1:100 dilution), EEA-1 (1:50 dilution), or a cocktail of primary antibodies at dilutions above.

Images were acquired using a Nikon A1R confocal microscope (equipped with 405 nm/Argon/561 nm lasers) using a Plan APO VC 60X/1.40 NA oil immersion objective with the Nikon Elements software (Nikon Instruments Inc.) or the Zeiss LSM 800 confocal microscope (equipped with 405 nm/488nm/561/640 nm lasers) using a Plan APO VC 63X/1.40 NA oil immersion objective with Zen 5 software (Zeiss). The X-Z and Y-Z cross-sections were produced by orthogonal reconstructions from z-stack scanning at 0.25 µm intervals taken with 60X/63X objective in a 5 µm thick paraffin-embedded tissue section or Caco-2 monolayers. Three-dimensional reconstructions were performed using the Nikon Elements (Nikon Instruments Inc.) or the Zen 5 software. Post-acquisition processing was done in Adobe Photoshop.

To enumerate the number of infected cell types, the number of *L. monocytogenes* positive cells from each of the following cell types: a) extruding cells, (b) goblet cells, and (c) other intestinal epithelial cells (IEC’s) were counted from randomly acquired immunostained images from 45 intact villi (15 villi each from 3 gerbil/ treatment) and expressed as a number of infected cells/15 villi. For analysis of redistribution of cell–cell junctional proteins, randomly acquired images of Caco-2 cells or mouse ileum from five different fields from three independent experiments (for Caco-2 cells, representing 90–100 cells) or three individual mice (representing 100–150 epithelial cells) per treatment were acquired. The relative expression levels of MLCK and P-MLC were analyzed using the NIH ImageJ software. To analyze the redistribution of cell– cell junctional protein, the total number of cells containing intracellular cell–cell junctional protein puncta were manually counted in randomly acquired images and calculated, respectively.

### Analysis of *in vivo* intestinal permeability

Gerbils or mice were orally gavaged with 100 μl of non-metabolizable 4 kDa FITC-labeled dextran (FD4; 36 mg/100 µl (gerbils), of FD4; 15 mg/100 μl (mouse); Sigma Sigma-Aldrich) 4-5 h before sacrifice. Serum (50 µl each) collected above was mixed with an equal volume of PBS, and fluorescence was measured (Ex: 485 nm; Em: 520 nm; Spectramax, Molecular Devices), and the FD4 concentration was calculated using a standard curve generated by serially diluting FD4 in PBS. The serum from the uninfected gerbil or mice determined the control levels.

### Epithelial Permeability, Bacterial Translocation, Invasion, Pharmacological Inhibitors and Co-infection

Caco-2 cells were grown as monolayers on Transwell inserts with 3.0 mm pores (Corning-Costar) for up to 14-21 days. TEER was measured to monitor the monolayer integrity (Millicells Voltmeter, Millipore). A TEER value of at least 200 U/cm^2^ (±10) was used as the basal value to monitor the monolayer integrity (44). Bacterial cells were washed three times in PBS and resuspended in DMEM-FBS (10%) at an MOI of 50 and added to the Transwell system’s apical side. After 2 h incubation period at 37°C in 5% CO_2_, the liquid was collected from the basal well. Translocated bacteria were enumerated by plating (44).

For bacterial invasion analysis, monolayers were washed with PBS after 1 h of infection (MOI, ∼50) and incubated with DMEMFBS (10%) containing gentamicin (50 mg/ml) for 1 h. Caco-2 cells were lysed with 0.1% Triton X, and the internalized bacteria were enumerated by plating.

For analysis of FD4 flux, non-metabolizable 4 kDa FITC-labeled dextran (FD4; 5 mg/ml, Sigma-Aldrich) was added with bacteria (MOI, 50) resuspended in DMEM-FBS (10%), and added to the apical side. After 2 h incubation at 37C in 5% CO_2_, the liquid was collected from the basal well, and fluorescence was measured (Ex: 485 nm; Em: 520 nm; Spectramax, Molecular Devices).

For pharmacological inhibition of endocytic pathways, Caco-2 cells were pretreated with the macropinocytosis inhibitor amiloride hydrochloride (50 µM, Sigma) or chlorpromazine (CPZ; 10 µg/ml, Sigma) and pitstop-2 (50 µM, Sigma), which inhibit endocytosis via clathrin-coated pits, or methyl-β-cyclodextrin (MβCD; 5 mM) that inhibits caveolae-like membrane domains or the synthetic glycosphingolipid L-t-LacCer (β-d-lactosyl-N-octanoyl-l-threo-sphingosine; 5 µM, Avanti Polar Lipids) that blocks caveolar endocytosis without some of the off-target effects of MβCD (37) or the dynamin inhibitor dynasore (80 µM, Sigma) for 30 min prior.

For coinfection, the WT (Em^R^) and Lin^InlA(EmS)^ or *lap^─^*^(EmR)^ and Lin^InlA(EmS)^ were mixed at a 1:1 ratio (∼MOI 50 of each strain). For invasion assays, wells were washed thrice with serum-free DMEM and lysed with 1 ml of sterile 0.1% Triton X-100. For translocation assays, the liquid was collected from the basal well. Samples were serially diluted prior to plating on BHI agar (BHIA) and BHIA supplemented with 5 μg/ml of erythromycin. The number of colonies on BHIA plates without antibiotic (erythromycin) was subtracted from those on BHIA plates with 5 μg/ml of erythromycin. The results were expressed as % percentages of bacterial colonies invaded or translocated to mammalian cells.

### Western Blotting

*Listeria* strains were grown as above to assess the expression of LAP or InlA. To isolate cell wall-associated proteins, washed bacterial pellets from 10 ml of overnight-grown cultures were resuspended in 0.5 ml protein extraction buffer (0.5% SDS, 10 mM Tris at pH 6.9) and incubated at 37°C for 30 min with agitation. Halt proteases and phosphatase inhibitors (Thermo Fisher Scientific) were used during all protein extraction procedures. The protein concentrations were determined by BCA assay (Thermo Fisher Scientific) and separated on SDS-PAGE gels (10–12.5% polyacrylamide) and electro-transferred to the polyvinylidene difluoride (PVDF) membrane (Millipore). The membranes were then blocked in 5% nonfat dry milk (NFDM) in 0.1M Tris-buffered saline, pH 7.5 (TBS), containing 0.1% Tween 20 (TBST) for at least 1 h. All primary antibodies were diluted in 5% NFDM in TBST and incubated overnight. Primary antibodies (Table S2) against LAP or InlA (1 μg/ml). The HRP-linked secondary antibodies, anti-rabbit IgG or anti-mouse IgG (1:2000 dilution in 5% NFDM in TBST) were incubated at 37°C for 1 h, and a chemiluminescence method was performed using LumiGLO reagent (Cell Signaling) before visualization in the Chemi-Doc XRS system (Bio-Rad). To immunoprobe the same membrane with another antibody, the originally bound antibodies were stripped using Restore Western Blot Stripping Buffer (Thermo Fisher Scientific) according to the manufacturer’s protocol.

### Recombinant Protein Purification

Recombinant proteins (LAP) containing endogenous His, S, and Trx tags derived from the pET-32a/pET28 cloning vector (Novagen) from ClearColi (Lucigen) were purified using a Ni-affinity column. In ClearColi, LPS lacks a secondary acyl chain, thus eliminating endotoxicity. Briefly, for LAP purification, ClearColi were each grown in 1 L LB broth (BD) with ampicillin (50 mg/ml) for 3 hr at 37C and IPTG (1 mM) induced at 20C for 12 hr. After sonication (total 7 min, with cycles of 30 s sonication and 15 s pulse; Branson Sonifier), the supernatants were purified by Ni-column. Protein concentrations were measured by Bradford assay, and the purity was monitored by SDS-PAGE (10%-acrylamide). Purified recombinant InlA was provided by Marcelo Mendonca, (University of Pelotas, Brazil).

For re-association of externally added InlA to *Lin*, bacterial cells were harvested from 1 mL of overnight grown culture, and the pellet was washed three times in PBS before adding 1 or 2 µg/ml of purified LAP. The mixture was incubated for 30 min at 30C with continuous shaking and then pelleted, washed five times in the PBS, resuspended in DMEM, and used in the invasion as mentioned above or translocation assay.

### Statistical Analysis

Experimental data were analyzed using GraphPad Prism (La Jolla, CA) software. The Mann-Whitney test assessed statistical significance for gerbil or mouse microbial counts. In other experiments, comparisons between two data sets were performed using the unpaired Student *t*-test. When comparisons between more than two data sets were performed, a one-way or two-way analysis of variance with Tukey’s multiple-comparison test was performed. All data are representative of at least 3 independent experiments, and specific numbers of gerbils or mice per group are noted in corresponding figure legends. Unless otherwise indicated, data for all experiments are presented as the mean ± standard error of the mean (SEM).

## DATA AVAILABILITY

All strains used in this study will be made publicly available upon publication and upon request.

## ACKNOWLEDGMENTS

The research was supported by funds from the National Institute of Food and Agriculture of USDA (Hatch grant # 1016249), the KY-INBRE (NIH-sponsored, RD), and the start-up funds at EKU and ODU (RD). The authors acknowledge J. A. Schaber for assistance with confocal microscopy and G. Shafer and V. Bernal-Crespo for histopathology.

## AUTHOR CONTRIBUTIONS

RD and AKB generated the idea, designed the experiments, performed the experiments, interpreted the data, wrote the manuscript, and acquired funding. All other authors (ST, DBB, JT, BA, DL, NLFG, KKM, MS, MRS, DKM) assisted in conducting experiments. ACD helped with histopathology scoring. All authors reviewed the manuscript.

## SUPPLEMENTAL MATERIALS

(Table S1, S2 and Fig S1-S9)

